# Constitutive ATR Signaling Enforces Domain-Wide Origin Synchrony

**DOI:** 10.64898/2026.10.19.760027

**Authors:** Joyce Kamar, Sharon Eden, Sunil K. Pradhan, Maruthi K. Pabba, Maria Arroyo, Avraham Greenberg, Leonor Daniel, M. Cristina Cardoso, Itamar Simon

## Abstract

Eukaryotic DNA replication relies on organized spatial-temporal domains, yet how neighboring origins are coordinated at the single-molecule level remains unresolved. Combining single-molecule DNA combing with super-resolution microscopy, we demonstrate that during unperturbed early S-phase, adjacent replication origins across diverse vertebrate species fire synchronously as a coordinated unit. This domain-wide temporal harmony is continuously enforced by a baseline, constitutive ATR–CHK1 signaling axis that restrains local CDK2 and CDC7 kinase activities. Strikingly, replication stress dismantles this coordination, shifting origin activation toward an asynchronous, replicon-autonomous mode. Mechanistically, stress-induced phosphorylation of the cohesin subunit SMC1A promotes an SMC1A–ATR complex that sequesters ATR at stalled replication forks. This localized sequestration creates a regional ATR loss-of-function phenotype, relieving CDK2/CDC7 suppression and driving premature, uncoordinated origin firing. Super-resolution imaging shows that ATR sequestration structurally reorganizes replication foci, converting coordinated clustered foci into fragmented, lower-density clusters. Together, our findings uncover a baseline role for constitutive ATR signaling in enforcing domain synchrony and reveal a molecular switch that dynamically decouples origin activation during genomic stress to safeguard chromosome duplication.

## Introduction

Eukaryotic DNA replication is governed by a strict hierarchy of structural and temporal organization, spanning from individual genomic sites to large-scale nuclear compartments. At the foundation of this hierarchy are replication origins. Unlike in prokaryotes, eukaryotic origins are degenerate, scattered, and characterized by a lack of strict sequence specificity ^1^. When an origin fires, it activates a bidirectional replicon. Rather than operating in isolation, these replicons are organized into higher-order chromosomal structures known as replication domains or replication clusters, which loosely correspond to chromatin compartments mapped by Hi-C ^2^. Within the 3D nuclear space, classical fluorescence microscopy identifies these structural domains as replication foci (RFi) ^3,4^. Recent advances in super-resolution microscopy have further fragmented this architecture, revealing that individual macro-RFi are comprised of smaller sub-resolution clusters termed nano-foci (nRFi), which represent physically clustered, actively operating replicons ^5,6^.

The overarching temporal schedule of this structural network is captured macroscopically by replication timing (RT) maps ^7–9^. These population-average genomic profiles divide the chromosome into constant timing regions (CTRs), which represent unified replication domains that duplicate synchronously as a collective unit in either early or late S-phase, interconnected by timing transition regions (TTRs) where the replication schedule changes sequentially ^9–11^. RT programs are remarkably robust, evolutionarily conserved, and strictly associated with specific epigenetic states, transcription activity, and 3D genome architecture ^12^.

However, while extensive research has elucidated the macroscopic behaviors of replication domains, our understanding of how individual replicons are coordinated within a single domain remains heavily limited. Because traditional replication maps rely on population averages, they blur single-molecule dynamics. Consequently, whether adjacent origins within a replication domain fire simultaneously or sequentially on a single DNA molecule remains an open question.

Previous research has demonstrated that an active replication origin can interfere with the activation of its neighbors; as a primary fork sweeps through the locus, it passively replicates adjacent dormant origins, enforcing a characteristic physical spacing between active initiation events ^13^. Conversely, a sequential “domino model” has been proposed to describe how an active origin triggers the subsequent replication of neighboring regions. This orderly, step-wise progression has been documented both at the macroscopic level between adjacent replication domains (CTRs) ^14–16^ and at the single-molecule level within timing transition regions (TTRs). Within these transition zones, gradual genomic duplication is driven either by the long-range progression of a single, highly processive replication fork ^10^ or by the sequential, wave-like activation of individual origins spaced along the locus ^11^.

However, whether two adjacent active origins within a single replication domain (CTR) fire in a coordinated way or are activated independently remains elusive. This uncertainty touches a foundational question in chromosome biology – namely, what constitutes the fundamental regulatory unit of replication activation: an isolated individual replicon or the entire replication domain? If the replicon operates as the primary autonomous unit, individual origins should activate independently or sequentially in a localized domino cascade. Conversely, if the replication domain itself functions as the unified activation unit, all constituent origins within that domain should fire simultaneously.

This fundamental question has been only partially addressed by previous studies, yielding conflicting paradigms. Early single-molecule autoradiography, limited by a broad 30-minute temporal resolution, suggested that neighboring origins generally fire synchronously, though significant outliers were observed ^17^. More recently, high-resolution DNA combing restricted to the human 14q11.2 locus challenged this view, indicating that adjacent origins within this specific region do not activate at the exact same moment ^13^. However, these early reports were strictly limited in their genomic scope, throughput, and physiological context, highlighting a critical need for a comprehensive, genome-wide investigation to fully map the temporal landscape of origin coordination.

When cells encounter exogenous or endogenous challenges to replication, collectively termed replication stress, the replication program undergoes spatial and kinetic rewiring ^18^. At the single-molecule level, replication stress slows down or stalls ongoing replication forks, resulting in a marked reduction in the overall fork rate (FR). To prevent complete replication collapse and ensure the genome is fully duplicated, cells activate a compensatory mechanism – dormant origin activation, which significantly shortens the inter-origin distance (IOD) by awakening nearby backup origins ^19,20^. In spite of these local changes in FR and IOD, the overall replication organization, captured in RT remain stable and is largely not affected by replication stress ^21^. How replication stress affects the order of origin activation within a replication domain remains entirely unknown.

Central to the fidelity of the replication program is the master checkpoint kinase ATR (Ataxia Telangiectasia and Rad3-related), which handles the dual responsibility of managing replication stress and monitoring normal S-phase progression (reviewed in ^22^). Under conditions of replication stress, the uncoupling of active helicases from stalled polymerases generates extended tracts of single-stranded DNA (ssDNA). This ssDNA is rapidly coated by Replication Protein A (RPA), creating a platform that recruits and hyper-activates ATR ^23^. Once activated, ATR orchestrates a robust global checkpoint response to stabilize stalled forks, prevent replisome collapse, and transiently suppress the firing of late-replicating origins across the genome ^24,25^. However, emerging evidence has changed the classic view of ATR as a dormant emergency brake, revealing that it also maintains a continuous, low-level constitutive activity during an unperturbed S-phase ^26–34^. This baseline ATR signaling acts as an essential molecular rheostat in healthy, growing cells, continuously fine-tuning replication fork velocity and preventing premature or ectopic origin firing by keeping the activities of CDK2 and CDC7 strictly bounded ^28,29,35,36^. While this baseline ATR-CDK2/Cdc7 axis is known to ensure replication organization in untreated conditions, how this constitutive signaling is localized to maintain temporal harmony between immediately adjacent origins remains to be elucidated.

Beyond the soluble enzymatic cascades that govern S-phase, the spatial-temporal program of replication is intimately regulated by structural components of the chromosome. Among these is the cohesin complex, which is actively loaded onto the genome at active replication forks, where they physically encircle the sister chromatids to mediate cohesion ^37^. Upon replication stress SMC1A (Structural Maintenance of Chromosomes 1), a core component of the cohesin complex, is phosphorylated by the master kinase ATR at dual conserved residues (Ser957 and Ser966) ^38^. Strikingly, this post-translational modification creates a critical molecular nexus: once phosphorylated, SMC1A binds ATR, forming a tightly regulated structural-enzymatic feedback loop ^39^. While this phosphorylation-dependent recruitment of ATR by SMC1A is known to be required for intra-S-phase checkpoint execution, how this physical interaction behaves under localized stress to influence the coordination of nearby, unchallenged replication origins has yet to be explored.

Despite the profound realization that eukaryotic chromosomes are duplicated within organized spatial-temporal domains, a fundamental knowledge gap remains regarding how individual replicons are coordinated at the single-molecule level, and whether this coordination adapts dynamically to varying environmental conditions. To address this question, we systematically investigated the temporal coordination of adjacent origins across the entire genome under both unperturbed and replication stress conditions. By combining single-molecule DNA combing with super-resolution microscopy, we demonstrate that during an unperturbed early S-phase, adjacent replication origins possess an evolutionarily conserved tendency to fire simultaneously as a synchronized unit, a phenomenon strictly enforced by a baseline, constitutive ATR-CDK2/CDC7 signaling axis. Strikingly, we show that this temporal harmony is dynamically dismantled under replication stress. Mechanistically, we reveal that replication stress triggers a complex between SMC1A and ATR, effectively trapping the kinase at stalled replication forks. This phosphorylation-dependent sequestration creates a localized ATR “loss-of-function” phenotype, lifting the continuous checkpoint-mediated inhibition on regional CDK2 and CDC7 activities and driving the uncoordinated, asynchronous activation of adjacent neighbors. Together, our findings redefine the baseline architecture of the replication program and uncover a novel mechanism that allows cells to dynamically safeguard chromosome duplication during genomic crises.

## Results

### Simultaneous Firing of Adjacent Origins is more prominent in Early S-Phase Replication Domains and is disrupted by replication stress

To understand the temporal coordination of DNA replication at a single-molecule level, we sought to determine the relative activation timing of adjacent origins on individual DNA strands. While origins are known to cluster within distinct replication domains, it remains unclear whether immediately adjacent origins within a cluster fire synchronously or independently. To address this, we developed a quantitative metric termed Relative Origin Activation Timing (ROAT), utilizing high-resolution DNA combing, which is built on a previously published approach^13,17^. Cells from three phylogenetically distinct lines, mouse embryonic fibroblasts (MEFs), human embryonic kidney cells (HEK293), and chicken DT40 cells, were sequentially pulse-labeled with IdU and CldU, followed by a 1.5-hour chase. Harvested DNA was combed onto silanized coverslips and immunolabeled for IdU (red), CldU (yellow), and single-stranded DNA (green) (**Figure 1a**). Individual replication origins were identified by characteristic symmetrical track patterns (CldU-IdU-unlabeled DNA (optional) –IdU-CldU). The size of the replication bulb, namely the distance between two diverting CldU tracks, serves as a direct proxy for its initiation timing relative to the CldU pulse: a larger bulb denotes an origin that fires long time before the CldU pulse, whereas a smaller bulb indicates an origin that initiated closer to the CldU pulse. Because bulb size is inherently confounded by local fork velocity, we normalized this measurement by dividing the bulb length by twice the average fork rate (FR) calculated for that specific slide:

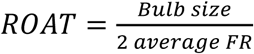

**Figure 1.**
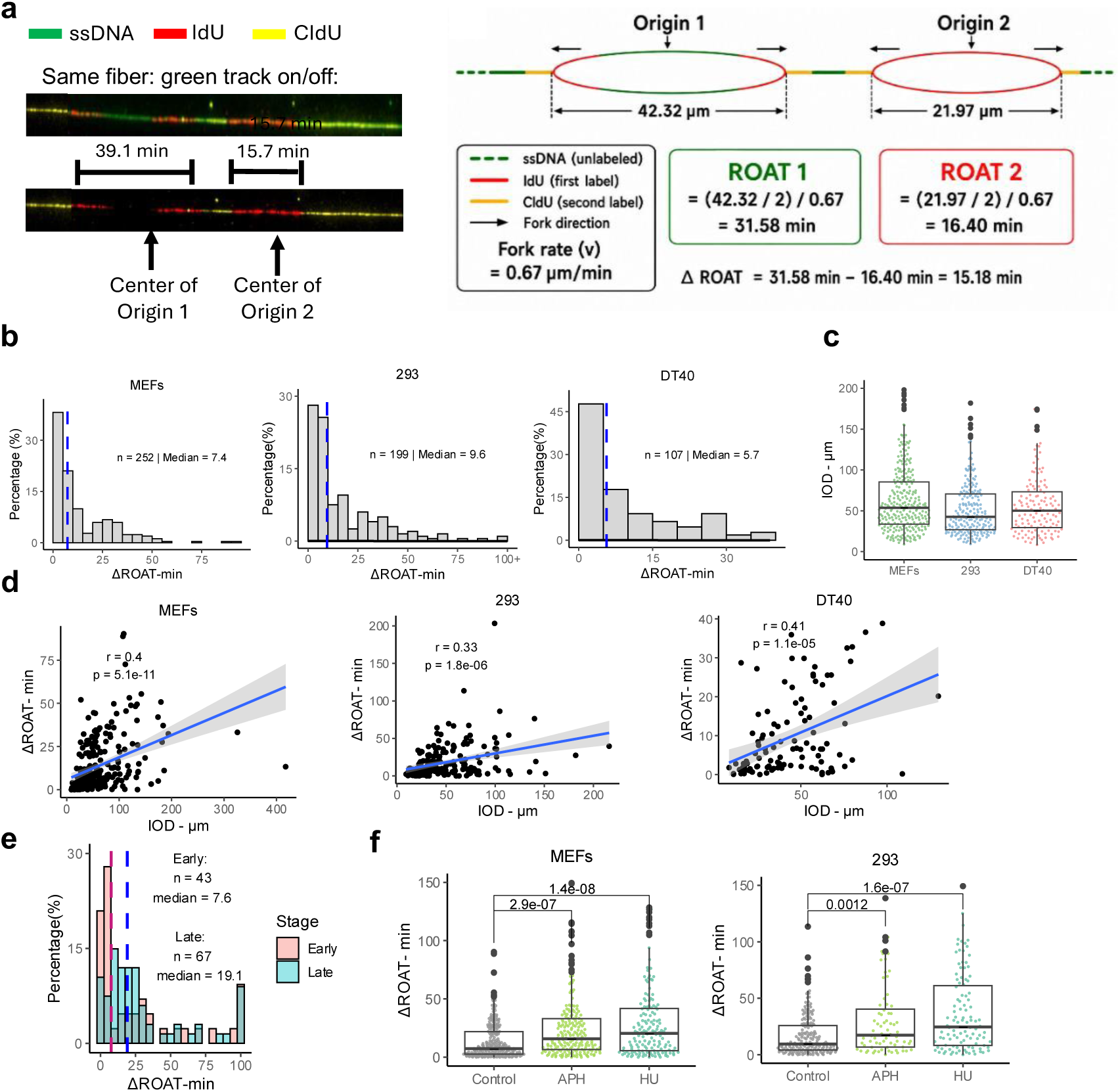
Adjacent replication origins display coordinated activation that is disrupted by replication stress. **(a)** left: representative DNA fiber, the same DNA fiber is shown with all channels (up) and without the ssDNA channel (down) to facilitate visualization of the IdU and CldU labeling patterns. The centers of the two adjacent origins and their corresponding ROAT values are indicated. Right: schematic representation of the calculation of relative origin activation timing (ROAT) and ΔROAT between two adjacent replication origins. **(b)** Distribution of ΔROAT values between adjacent replication origins in MEFs, HEK293, and DT40 cells. Dashed vertical lines indicate the median ΔROAT. The number of analyzed origin pairs (*n*) and median ΔROAT are indicated for each cell line. Normality of the ΔROAT distributions was assessed using the Shapiro–Wilk test (MEFs: W = 0.795*, p = <2.2*10^-16*; 293: W = 0.678*, p = <2.2*10^-16*; DT40: W = 0.826*, p = 6.65*10^-10*). **(c)** Inter-origin distance (IOD) measured between adjacent replication origins in MEFs, HEK293, and DT40 cells. Each point represents an individual IOD measurement. Box plots show the median (center line), first and third quartiles (box), and values extending up to 1.5× the interquartile range (whiskers). **(d)** Scatter plots capturing the correlation between IOD and ΔROAT in MEFs, HEK293, and DT40 cells. Each point represents an individual pair of adjacent origins. The regression line and corresponding confidence interval are shown. Pearson correlation coefficients (*r*) and corresponding *p* values are indicated in each graph. **(e)** Distribution of ΔROAT values in early– and late-S phase HEK293 cells. Cells were separated by FACS according to DNA content, and early– and late-S-phase fractions were collected and analyzed separately. Dashed vertical lines indicate the median ΔROAT for each population; the number of analyzed origin pairs (*n*) and median values are indicated. Statistical significance between early– and late-S-phase ΔROAT distributions was assessed using the Kolmogorov–Smirnov test (D= 0.332 p = 0.004). **(f)** ΔROAT between adjacent replication origins in untreated control cells and following replication stress induced by aphidicolin (APH) or hydroxyurea (HU) in MEFs and HEK293 cells. Each point represents an individual pair of adjacent origins. *P* values for the indicated comparisons (two-tailed t-test) are shown above the plots

For example, a replication bulb measuring 42.32 µm on a slide with a mean FR of 0.67 µm/min corresponds to an origin that fired 31.58 minutes prior to the CldU pulse (**Figure 1a**). This allowed us to calculate the precise temporal delta between neighboring initiation events on the same molecule

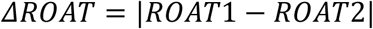

To evaluate the temporal coordination of neighboring replication events, we plotted the distribution of ΔROAT values across the three species (**Figure 1b**). In all three systems, the ΔROAT profiles of adjacent origins were heavily skewed toward the smallest temporal bins (0-10 minutes) and was significantly smaller than the ΔROAT between random pairs (**Figure S1a**), demonstrating an evolutionarily conserved tendency for adjacent origins to fire simultaneously. Notably, this synchronous activation was not absolute; approximately 44% of adjacent origin pairs exhibited a ΔROAT greater than 10 minutes (**Figure 1b**), indicating a distinct subpopulation of asynchronously firing neighbors.

To determine whether local structural or kinetic features influence this synchrony, we measured for each cell the fork rate (FR), the distance between two adjacent origins (IOD) and fork asymmetry, the latter serving as a marker for localized DNA damage^40^ (**Figures 1c and S1b-c**). While ΔROAT correlated with inter-origin distance (IOD), no correlation was observed with fork rate (FR) or fork asymmetry (**Figures 1d and S1b-c**). These findings suggest that origin firing coordination is independent of local fork velocity and localized DNA damage, but is modestly influenced by the linear spacing between adjacent origins. It should be noted that mammalian replication domains typically span several hundred kilobases to megabases ^9^, whereas the typical IOD observed in our data is <200 Kb (**Figure 1c**). Thus, the vast majority of the measured ΔROAT are expected to lie predominantly within a single replication domain and thus reflects the coordinated activation of adjacent origins within a domain.

Next, we investigated whether this simultaneous firing phenomenon is uniform throughout the S phase. We used FACS to isolate HEK293 cells in early and late S-phase, and calculated the ΔROAT for each fraction (**Figure 1e**). Strikingly, origin firing synchrony exhibited a strong dependence on S-phase progression. While adjacent origins in early S-phase fired in tight temporal alignment, late S-phase populations displayed a significantly broader ΔROAT distribution, with the median temporal delta 2.5-fold higher than in early S-phase (**Figure 1e**). Nevertheless, late-replicating adjacent origins still initiated within a relatively constrained temporal window (median ΔROAT ∼19 minutes), well below the average time required to complete full replicon duplication. Taken together, these results demonstrate that tight, simultaneous origin firing is a prominent hallmark of early S-phase replication, whereas late S-phase transitions toward a less synchronized, more temporally relaxed mode of origin activation that remains largely independent of local fork kinetics.

To further characterize simultaneous firing of adjacent origins we explored how this coordinated timing program responds to replication stress. To test this, we exposed MEF and HEK293, cells to low-dose replication stress, using either the DNA polymerase inhibitor aphidicolin or the ribonucleotide reductase inhibitor hydroxyurea. Strikingly, induction of replication stress (observed by the reduction of FR and IOD; **Figure S1d-e**) significantly increased the ΔROAT of adjacent origins in both cell lines (**Figure 1f**), without changing S phase distribution (**Figure S1f**). This indicated that the baseline mechanism enforcing simultaneous firing is highly sensitive to, and dismantled by, replication stress.

### Baseline ATR Signaling Imposes the Synchronous Firing of Adjacent Origins

Having demonstrated that replication stress actively dismantles the simultaneous firing of adjacent origins, we next sought to identify the molecular machinery governing this temporal coordination under normal growth conditions. We focused our attention on the master checkpoint kinase ATR. Although ATR is traditionally characterized as a stress-activated responder to stalled replication forks, growing evidence indicates that it also maintains a low level of baseline, constitutive activity during an unperturbed S-phase to regulate normal replication fork progression and suppress global late-origin firing ^26–33^. We hypothesized that this baseline ATR signaling might be actively required to maintain the synchronous activation of neighboring origins in unstressed cells. To test this hypothesis, we chemically inhibited ATR in untreated, actively growing cells using a panel of structurally distinct ATR inhibitors, including VE-821, VE-822, AZD6738, under normal growth conditions. Intriguingly, treating unstressed cells with these ATR inhibitors (ATRi) completely recapitulated the replication stress phenotype, significantly increasing ΔROAT and disrupting the simultaneous firing of adjacent origins (**Figure 2a**). To rule out potential off-target effects of chemical inhibition, we utilized DT40 cells harboring an auxin-inducible degron (AID)-tagged ATR ^41^. Genetic degradation of ATR yielded similar results, confirming that both acute chemical inhibition and targeted degradation of ATR lead to a severe loss of origin coordination (**Figure 2a**). Importantly, this phenotype was highly specific to the ATR pathway; chemical inhibition of the related kinase ATM had no discernible effect on ΔROAT, despite exerting a similar impact on baseline fork rate (FR) and inter-origin distance (IOD) (**Figures 2a and S2a-e**). These data reveal a novel, baseline requirement for ongoing ATR activity, but not ATM, in maintaining synchronous origin activation under normal growth conditions.

**Figure 2.**
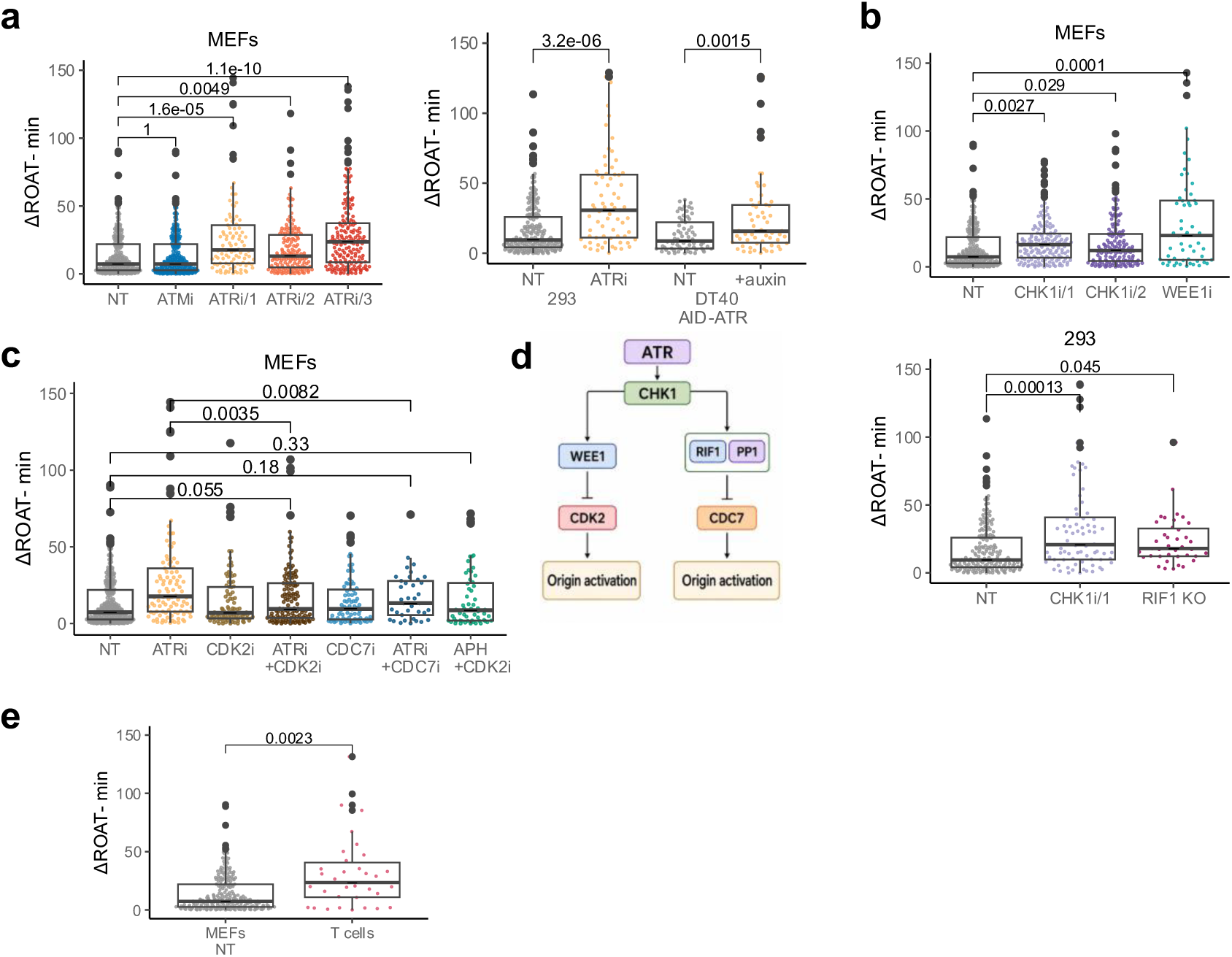
ATR–CHK1 signaling maintains coordination between adjacent replication origins through regulation of CDK2 and CDC7. **(a)** ΔROAT between adjacent replication origins in MEFs (left) and HEK293 cells (right), was measured in untreated (NT), ATM inhibition (ATMi) or ATR inhibition using three different ATR inhibitors (**methods**). ΔROAT was also measured in DT40 AID-ATR cells in the absence or presence of auxin-induced ATR depletion. **(b)** ΔROAT measurements in MEFs and HEK293 cells. MEFs were analyzed under untreated (NT), CHK1 inhibitor-treated (CHK1i), and WEE1 inhibitor-treated (WEE1i) conditions (upper pannel), while HEK293 cells were analyzed under untreated (NT), CHK1 inhibitor-treated (CHK1i), and RIF1 knockout (RIF1 KO) conditions (lower pannel). **(c)** ΔROAT measurements in MEFS either untreated (NT), or with each of the following: ATRi(VE-821), CDK2i, ATRi(VE-821) together with CDK2i, CDC7i, ATRi(VE-821) together with CDC7i or APH together with CDK2i. **(d)** Proposed model for the regulation of replication-origin activation by the ATR– CHK1 pathway. **(e)** ΔROAT measurements in untreated MEFs (NT) and activated primary CD8+ T cells. See **Methods** for details of splenocyte activation and CD8+ selection. *P* values for the indicated comparisons (two-tailed t-test) are shown above the plots (a-c,e). Details about the inhibitors are in the Methods section.

To further map the downstream ATR signaling cascade responsible for regulating simultaneous origin firing, we evaluated the effects of targeting its classical downstream effectors. Similar to direct ATR disruption, chemical inhibition of CHK1 and WEE1 as well as *Rif1* KO led to a corresponding increase in ΔROAT (**Figure 2b, and S2f-i**). Because ATR classically restricts premature origin firing by suppressing Cyclin-Dependent Kinase 2 (CDK2) and CDC7 activity ^28,29,35,36^, we next examined whether deregulated CDK2 and CDC7 signaling was responsible for the uncoordinated firing phenotype observed upon ATR disruption. Remarkably, co-treatment of cells with ATRi and either a selective CDK2 inhibitor or CDC7i rescued the ATRi-induced defect, restoring the low ΔROAT distribution characteristic of synchronous adjacent origin firing (**Figure 2c, and S2j-k**). These pharmacological rescues, observed with both CDK2i and CDC7i, are consistent with ATR constraining CDK2/CDC7 to maintain synchrony, strongly supporting the model that hyper-activation of dormant or premature origins underlies the uncoordinated firing of adjacent neighbors. Interestingly, stress induced synchronization loss was also CDK2 dependent and cell treated with aphidicolin in the presence of CDK2i retained synchronization (**Figure 2C and S2j-k**). Taken together, these results support a model in which baseline ATR maintains origin synchrony, at least in part by restraining CDK2 and CDC7 kinase activities (**Figure 2d**).

This requirement for low baseline CDK2 activity prompted us to ask whether physiological contexts characterized by naturally elevated CDK2 levels would likewise lack spatial-temporal origin coordination. To test this hypothesis, we examined activated primary CD8⁺ T cells, which express elevated endogenous levels of CDK2 during proliferation^42^. Consistent with our model, unperturbed activated CD8⁺ T cells failed to maintain origin synchrony between adjacent replicons (**Figure 2e**).

### SMC1A Sequesters ATR Under Replication Stress to Disrupt Local Origin Synchrony

Our finding that baseline ATR activity is required for synchronous origin firing raised a compelling paradox: why does replication stress, which robustly activates the ATR checkpoint globally, phenocopy a localized loss of ATR function at adjacent origins? To resolve this, we first examined the epistatic relationship between replication stress and checkpoint inactivation by measuring ΔROAT in cells subjected to a combined treatment of either aphidicolin (APH) or HU and an ATR inhibitor (ATRi). Concomitant treatment with both agents resulted in a non-additive change in origin coordination compared to either treatment alone (**Figure 3a, and S3a-b**). This lack of synergy is consistent with an epistatic relationship between replication stress and ATR inactivation (although an assay saturation cannot be formally excluded). These findings suggest convergence on a shared pathway to disrupt origin synchrony. We therefore hypothesized that replication stress physically sequesters or “traps” ATR at stalled forks, spatially restricting its diffusion and preventing it from executing its constitutive, baseline suppressive functions on surrounding origins.

**Figure 3.**
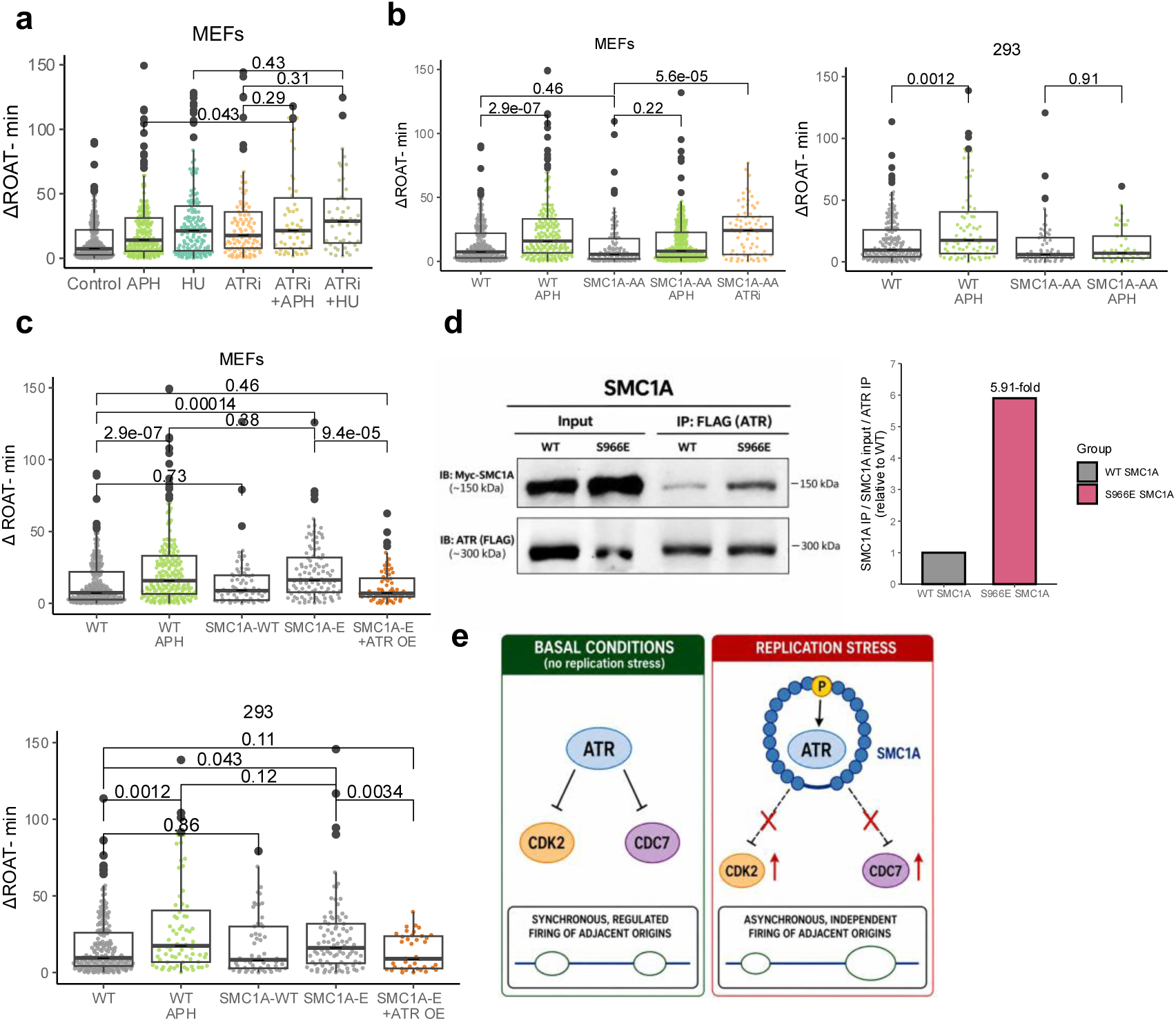
SMC1A contributes to the loss of coordinated replication-origin activation under replication stress through regulation of ATR signaling. **(a)** ΔROAT between adjacent replication origins in MEFs following replication stress and ATR inhibition. ΔROAT was measured in control, APH-, HU-, and ATRi-treated cells, and combined ATRi and APH or HU treatments. **(b)** ΔROAT measurements in WT and SMC1A-AA MEFs (left) and HEK293(right) cells with or without APH treatment in both cell lines, and with ATRi(VE-821) in MEFs. (**c)** ΔROAT measurements in WT, SMC1A –WT, SMC1A –E with or without ATR overexpression (OE) in MEFs (upper panel) and HEK293 cells (lower panel). *P* values for the indicated comparisons (two-tailed t-test) are shown above the plots (a-c). **(d)** Co-immunoprecipitation analysis of WT and S966E myc-tagged SMC1A with ATR. Representative immunoblots (left) showing myc-SMC1A and FLAG-ATR input, and after immunoprecipitation with FLAG-ATR. Quantification (right) of SMC1A association with ATR was calculated as the myc-SMC1A IP signal normalized to myc-SMC1A input and ATR IP, and expressed relative to WT SMC1A. (**e)** Proposed model for SMC1A –dependent regulation of origin coordination.

A primary candidate for this physical trapping mechanism is SMC1A, a core structural component of the cohesin complex with established roles in the intra-S-phase checkpoint. Upon replication stress, SMC1A is directly targeted and phosphorylated at dual conserved residues (Ser957 and Ser966) ^38,39^. To rigorously test whether this SMC1A-mediated sequestration of ATR is functionally required to dismantle origin coordination under stress, we utilized a genetic replacement strategy. We reasoned that if ATR trapping breaks synchrony, cells unable to form this complex would preserve origin coordination even during an active stress response. Indeed, mouse embryonic fibroblasts (MEFs) where endogenous Smc*1a* was replaced with the non-phosphorylatable *Smc1a*-AA mutant ^43^ fully preserved their origin coordination under aphidicolin stress, maintaining low ΔROAT values, indistinguishable from unstressed wild-type controls (**Figure 3b, and S3c-d**). Consistent results were observed upon the ectopic expression of *SMC1A*-AA in HEK293 cells (**Figure 3b, and S3e-f**). Conversely, we evaluated the consequences of forcing this trapping mechanism in the absence of stress by overexpressing a phospho-mimicking mutant of *SMC1A*, engineered for this study (*SMC1A*-E; **Methods**), which structurally models the stressed state constitutively. Strikingly, unperturbed cells expressing *SMC1A*-E (but not those expressing *SMC1A* WT) displayed a severe loss of origin coordination under baseline growth conditions, perfectly recapitulating the high ΔROAT phenotype observed during wild-type replication stress (**Figure 3c, and S3g-l**). To definitively confirm that this asynchronous firing is driven by a localized dilution of free, functional ATR rather than an active, dominant-negative signaling effect of the complex, we tested whether an excess pool of ATR could rescue the phenotype. Remarkably, the overexpression of wild-type ATR restored baseline origin synchrony and prevented coordination loss, even in the presence of the constitutive SMC1A –E mutant (**Figure 3c**). Furthermore, by performing Co-immunoprecipitation using in vitro–translated Myc–SMC1A and lysates from FLAG–ATR–expressing cells, we were able to show stronger association of ATR with phospho-mimicking SMC1A (S966E; **Figure 3d**). Taken together, these biochemical and genetic data support a model in which replication stress promotes a phosphorylation-dependent SMC1A–ATR complex that functionally sequesters ATR at stalled forks, preventing it from performing its baseline role of inhibiting regional CDK2 and CDC7 activities, which ultimately drives the asynchronous, independent activation of adjacent replication origins **(Figure 3e).**

### Super-Resolution Imaging of Replication Foci Architecture Validates the Loss of Origin Synchrony

While our single-molecule DNA combing analysis established a clear temporal coordination between neighboring origins, we sought to determine how this spatial-temporal organization manifests within intact nuclear architecture. To this end, we utilized correlative confocal and super-resolution microscopy to interrogate the structural composition of individual replication foci (RFi). By combining confocal imaging with super-resolution microscopy, we quantified the number of active, individual replicons (nano-RFi) contained within individual replication focus (RF) during a brief 10-minute EdU pulse (**Figure 4a; Methods**). This visual framework effectively expanded our observations from isolated adjacent origin pairs to the broader multi-origin collective of an entire replication focus.

**Figure 4:**
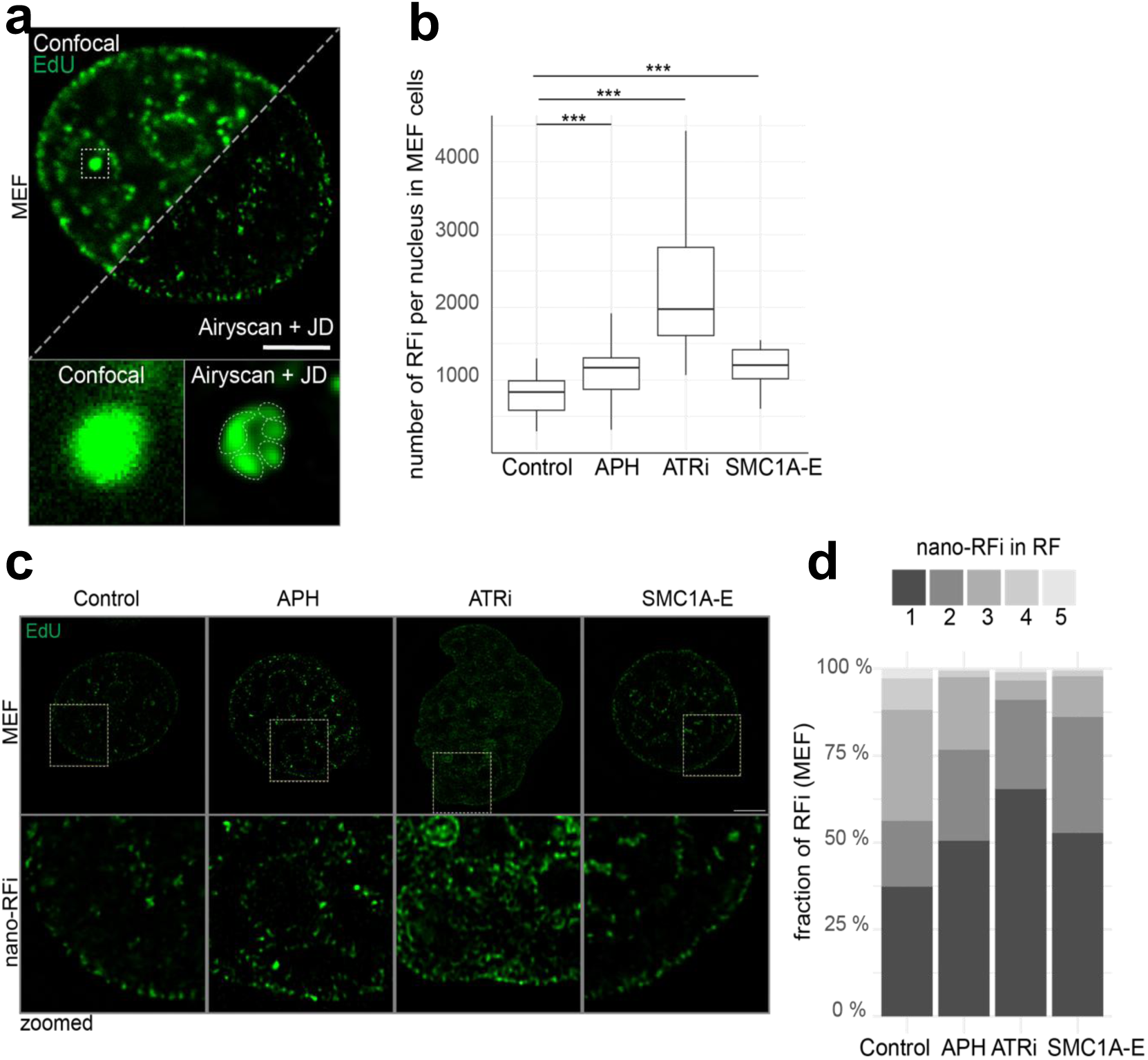
Loss of replicon coordination upon ATR inhibition. **(a)** Correlative microscopy of the same nucleus of mouse embryonic fibroblasts (MEF) shows replication foci at confocal (RFi) and super-resolution (nano-RFi) resolution. A single RFi at confocal resolution corresponds to a cluster of nano-RFi/replicons. Scale bar: 5 µm. **(b)** Boxplot of the number of RFi per nucleus across experimental conditions, (n = 73 nuclei; Control n = 18, Aphidicolin n = 23, ATRi n = 20, SMC1A-E n = 12). Boxes span the 25th–75th percentiles, the line depicts the median, and whiskers extend to 1.5 × IQR. Statistical significance relative to control was assessed using the Wilcoxon rank-sum test with Holm correction for multiple comparisons (n.s. for p ≥ 0.05; * for p < 0.05 and ≥ 0.005; ** for p < 0.005 and ≥ 0.0005; *** for p < 0.0005). p-values: Aph– control p = 2.2 x 10^-5^, ATRi–control p = 8.7 x 10^-7^, SMC1A-E–control p = 7 x 10^-5^. **(c)** Representative super-resolution (Airyscan + joint deconvolution) images of nano-RFi in each condition, with magnified regions shown below for better visualization (boxed areas in the upper panels). Scale bar: 5 µm. **(d)** Fraction of RFi containing 1, 2, 3, 4 or ≥ 5 nano-RFi, inferred from correlative microscopy. RFi and nano-RFi were segmented from confocal and super-resolution images of the same cells, and the number of nano-RFi was counted within each individual RFi.

Under normal, unperturbed growth conditions, early S-phase cells displayed a balanced distribution of well-defined replication foci, each containing tightly clustered nano-foci (**Figure 4a**). Strikingly, conditions that disrupted single-molecule origin coordination, including aphidicolin-induced replication stress (APH), ATR inhibition (ATRi), or expression of the novel phospho-mimicking SMC1A-E mutant, triggered a significant increase in the total number of replication foci per nucleus (**Figure 4b**). This systemic increase in active foci across the nucleus is presumably driven by the unchecked, premature activation of CDK2 and CDC7 kinase activities.

Crucially, this increase in global foci density per cell was accompanied by a sharp decrease in the internal density of active replicons per individual focus, reflected by a significantly reduced nano-RFi to macro-foci ratio (nRFi/RF; **Figure 4c,d**). This physical reorganization closely mirrors our single-molecule combing observations: when baseline ATR activity is suppressed or spatially sequestered, the systemic repression of CDK2 and CDC7 is lifted, allowing an aberrant number of distinct replication foci to prematurely initiate across the nucleus. However, because internal spatial-temporal coordination within each focus is broken, fewer constituent origins activate synchronously within a given domain, resulting in structurally fragmented, “emptier” foci characterized by a diminished nRFi/RF ratio.

Together, these super-resolution imaging data provide independent physical and visual validation of our molecular model. They demonstrate that the loss of origin coordination captured by the ΔROAT metric manifests within the nucleus as a structural reorganization of the replication machinery, wherein a greater number of distinct replication domains are activated globally, but the synchronous firing of neighboring origins within those domains is dismantled.

## Discussion

In this study, we have uncovered a fundamental regulatory mechanism governing the spatial-temporal program of eukaryotic DNA replication. We demonstrate that during an unperturbed early S-phase, adjacent replication origins possess an evolutionarily conserved tendency to fire simultaneously as a synchronized unit. This temporal coordination is strictly enforced by a baseline constitutive ATR-CHK1 signaling axis that dynamically restrains local CDK2 and CDC7 activities. Strikingly, we show that this domain-based synchrony is dynamically dismantled under replication stress. Mechanistically, our data are consistent with a model in which replication stress promotes a phosphorylation-dependent SMC1A–ATR complex that can sequester ATR at stalled forks. In fact, the level chromatin bound ATR was shown to increase in aphidicolin and hydroxyurea treated mouse embryonic stem cells ^44^ in agreement with it becoming trapped by SMC1, as shown here. This phosphorylation-dependent trapping creates a local “loss-of-function” phenotype for ATR, lifting the continuous inhibition on CDK2/CDC7 and driving the uncoordinated, asynchronous activation of adjacent neighbors. Future work is needed to directly quantify ATR redistribution in vivo under stress and SMC1A mutants.

### Methodological Advantages of the ΔROAT Metric

To accurately measure the activation kinetics of individual replicons in a manner that permits direct comparison across diverse physiological states, we developed the Relative Origin Activation Timing (ROAT) metric. Previously, the replication bulb size was used as a proxy for the relative time of activation of an origin ^13,17^. We added to it a normalization step in which the replication bulb size is divided by twice the average replication fork rate, calculated for that specific genomic environment. Crucially, this normalization enables a direct comparison of origin firing coordination between completely disparate growth conditions.

Our finding that synchronous replication of adjacent origins is more prominent in early S and to normal unperturbed growth conditions, reconcile previous observation that found significant degree of asynchrony ^13,17^. Moreover, by scaling our ΔROAT metric genome-wide, we overcame the statistical limitations of locus-specific architectures ^13^. Finally, the consistent observation of synchronous origin firing across phylogenetically diverse species, including human, mouse, and chicken, strongly highlights the evolutionary conservation of this regulatory architecture.

### Potential Methodological Confounders

Because both replication stress and ATR inhibition alter ΔROAT alongside basic replication parameters, such as inter-origin distance (IOD) and fork rate (FR), we considered whether the observed shifts in ΔROAT might simply represent downstream consequences of altered fork kinetics or origin spacing. Two independent lines of evidence argue strongly against this possibility. First, we identified several specific perturbations, namely ATM inhibition (ATMi; **Figure 2a and Figure S2a,b**) as well as CDK2 and CDC7 inhibition (CDK2i, CDC7i; **Figure 2c and Figure S2j,k**) that significantly reduce both FR and IOD without altering ΔROAT. Second, single-molecule correlation analyses revealed no association between FR and ΔROAT, and only a modest association with IOD (**Figure 1d and Figure S1a**). Together, these findings indicate that local fork velocity and inter-origin spacing do not serve as primary confounding variables driving origin firing synchrony.

Finally, because origin firing coordination is more pronounced during early S-phase, we evaluated whether the loss of synchrony observed under replication stress or ATR inhibition could be an indirect artifact stemming from shifts in S-phase cell cycle distribution. However, cell cycle profiling demonstrated that the overall S-phase population distribution remains comparable across these treatment conditions (**Figure S1f**), excluding S-phase staging artifacts as a driver of origin desynchronization.

### Domain-Wide Coordination vs. Isolated Adjacent Events

Because our DNA combing measurements are inherently limited to tracking the spatial-temporal relationship between two immediately adjacent active origins, they raise a fundamental question: is this strict temporal coordination restricted to isolated neighbor pairs, or does it reflect a global, domain-wide phenomenon? Logically, if every given pair of adjacent origins within a replication domain possesses a near-zero temporal delta (ΔROAT), this pairwise harmony must propagate across the entire locus. The absolute coordination of adjacent pairs that we observe therefore implies a synchronized, domain-wide execution of the replication program. The propagation of synchronization across an entire domain is strongly supported by early autoradiography work documenting simultaneous initiation within larger clusters of three to eight contiguous replicons ^17^. This single-molecule logic is visually and structurally confirmed by our super-resolution microscopy data. The sharp reduction in the nano-replication foci to macro-replication focus (nano-RFi/RF) ratio observed upon ATR inhibition, aphidicolin stress, or in SMC1A-E mutants provide direct physical evidence for a domain-wide regulatory model. Under normal baseline conditions, a continuous ATR signaling threshold keeps the entire replication focus operating as a single operational unit, forcing all constituent origins to fire in tandem. When this core regulatory axis is disrupted, a greater number of macro-replication foci across the nucleus are prematurely awakened due to unchecked global CDK2 and CDC7 activity, which is also clear from the super-resolution microscopy data. However, the internal synchronization within each individual focus is severely compromised. This structural breakdown transforms the unified replication focus into a “de-synchronized” cluster of isolated nano-RFi, where individual replicons fire in a scattered, uncoordinated sequence.

To explain how baseline ATR activity enforces domain-wide synchrony, we propose a “limited resource” model. During an unperturbed early S-phase, essential, rate-limiting replication initiation factors (such as TopBP1, MTBP, and pre-initiation complex components) exist in finite quantities and must be efficiently deployed across the nucleus. We speculate that baseline ATR activity acts as a molecular gatekeeper that tightly regulates access to these limited resources. By actively suppressing sporadic, ectopic origin firing at unlicensed or heterochromatic loci, ATR ensures that these finite firing factors are concentrated and selectively “tunneled” to specific, designated early replication domains. When a domain is licensed to open, this locally enriched pool of firing factors enables all internal origins to fire simultaneously in a coordinated burst.

Under replication stress, phosphorylation-dependent trapping of ATR by SMC1A at stalled forks creates a localized loss of ATR function at those specific loci. As numerous forks stall across the genome, this widespread physical sequestration functionally depletes the pool of free, nucleoplasmic ATR. Indeed, it was shown that under replication stress ATR moves from the nucleoplasm into discrete foci ^45^ and its chromatin bound fraction increases^44^. This systemic depletion or direct chemical inhibition via ATRi, lifts the global inhibition on CDK2 and CDC7. Consequently, an aberrant excess of replication domains across the nucleus is prematurely awakened. This global explosion of active foci scatters the finite pool of firing factors across the entire genome, starving individual foci of the threshold concentrations required for synchronized initiation. Thus, local ATR sequestration triggers a global resource-dilution cascade, transforming unified, highly synchronized macro-foci into fragmented clusters of scattered, asynchronous nano-foci.

### Replicon-Based vs. Domain-Based Organization: A Stress-Induced Switch

Our data reveal that the eukaryotic nucleus balances two competing topological and kinetic strategies for genome duplication: domain-based organization and replicon-based organization. Under physiological growth conditions, the cell treats the entire replication domain, visually represented by the macro-replication focus (RF), as a unified operational unit wherein all constituent origins fire together in a tightly synchronized burst. Intriguingly, this tight temporal coordination is most pronounced during early S-phase, gradually relaxing in late S-phase populations where origin firing adopts a less synchronized, more autonomous replicon-based pattern. We hypothesize that this temporal shift reflects a reorganization of cellular priorities as S-phase progresses. Early S-phase is heavily enriched for actively transcribed genes ^46^. Because active replication foci transiently suppress or compromise local transcription ^47^, there is a distinct evolutionary advantage in compressing the duration of active replication within a given domain. Synchronizing all constituent origins within an early domain minimizes the duration of local transcriptional silencing, thereby safeguarding transcriptional fidelity. In contrast, late-replicating regions are generally heterochromatic, gene-poor, and naturally exhibit higher baseline mutation frequencies. In these genomic territories, maintaining a rigid, resource-heavy domain coordination is less vital than ensuring that every remaining stretch of the chromosome is fully duplicated prior to mitosis. By shifting away from strict domain-wide synchrony toward a more flexible, replicon-autonomous framework in late S-phase, the cell maximizes its ability to complete genome duplication under tight temporal constraints.

Under replication stress, we observe a general loss of origin coordination, demonstrating that even early-replicating origins abandon domain-wide synchrony. When an ongoing replication fork encounters an obstacle and stalls, maintaining a blind, domain-wide firing program would be harmful: if all origins within a domain were to fire simultaneously into a damaged or dNTP-depleted environment, dozens of forks would stall concurrently, requiring an unmanageable amount of simultaneous repair and rescue machinery. Shifting to an autonomous, replicon-based framework under stress allows the cell to decouple neighboring origins, managing each stalled fork individually. Thus, during genomic crises, the regulatory balance shifts from preserving transcriptional timing to safeguarding absolute replication fidelity.

### Pathological Outcomes of Perturbing the Spatial-Temporal Coordination Switch

Based on these findings, we hypothesize that any failure to properly execute this spatial-temporal program, either by disrupting the domain-based synchrony required during early S-phase or by failing to switch to an autonomous replicon-based strategy during replication stress or late S-phase, is highly detrimental to genome stability. For instance, if cells are unable to trap ATR via SMC1A during replication stress, adjacent origins continue to fire synchronously into damaged or compromised genomic zones. This uncoordinated continuation of domain-wide firing leads to simultaneous multi-fork collapse and extensive double-strand break formation. Conversely, if domain-wide coordination is permanently broken during unperturbed growth (e.g., in SMC1-E mutants or under chronic ATR inhibition), the continuous, premature awakening of scattered replication domains across the nucleus prevents the concentrated, efficient duplication of early replication domains. Scattering limiting replication factors across premature foci starves active domains of essential factors, prolongs the lifespan of single-stranded DNA intermediates, and increases the frequency of transcription-replication collisions. Ultimately, the SMC1A-ATR-CDK2/CDC7 axis serves as a vital molecular rheostat, balancing the structural integrity and transcriptional alignment of an unperturbed S-phase with the plastic, adaptive demands required to survive genomic stress. Our newly characterized *SMC1A* mutants offer a unique genetic toolkit to dissect these distinct operational modes, setting the stage for future investigations to definitively unravel the pathological consequences of breaking this replication checkpoint switch.

## Methods

### Cell culture and treatments

Mouse embryonic fibroblasts (MEFs; ATCC, CRL-299) were cultured in Dulbecco’s Modified Eagle Medium (DMEM, Biological Industries (Sartorius)) supplemented with 20% fetal bovine serum (FBS), 1% penicillin–streptomycin (Pen/Strep), and 1% glutamine. HEK293 cells (ATCC, CRL-1573) were cultured in DMEM supplemented with 10% FBS, 1% Pen/Strep, and 1% glutamine. MEFs and HEK293 cells were maintained at 37°C in a humidified incubator containing 5% CO₂.

For the CD8+ T cells in Figure 2f, cells were surplus from other experiments being performed in our lab, with all appropriate care being taken for the welfare of the mice prior to our experiment. Mice were C57BL/6J wild-type mice (The Jackson Laboratory, strain 000664). The spleen was collected and processed to obtain splenocyte suspensions. Red blood cells were lysed using Gibco red blood cell lysis buffer (Gibco A10492-01) according to the manufacturer’s instructions. Cells were subsequently washed and counted using a DeNovix CellDrop BF brightfield cell counter. Mouse T cells were activated using purified anti-CD3ε antibody (clone 145-2C11, BioLegend, at 1 µg/ml) and anti-CD28 antibody (clone 37.51, BioLegend, at 0.5 µg/ml). Activated splenocytes were maintained in RPMI (Sigma Aldrich, 11875093) containing 10% Fetal Bovine Serum (Gibco), 2 mM Glutamine (Satorius, 03-042-1B), 100 U/ml penicillin-streptomycin (Satorius, 03-031-1B and 50 uM Beta-mercaptoethanol (β-ME). CD8+ cells were positively selected using the CD8a (Ly-2) MicroBeads (Miltenyi Biotec) kit, following the manufacturer’s instructions.

AID-tagged ATR DT40 cells were kindly provided by Prof. Noel Francis Lowndes (Genome Stability Laboratory, Centre for Chromosome Biology, National University of Ireland Galway, Ireland) and were cultured in RPMI medium (Biological Industries (Sartorius)) supplemented with 20% FBS, 1% chicken serum, 1% Pen/Strep, 1% glutamine, and 10 mM HEPES. Cells were maintained at 39.5°C in a humidified incubator containing 5% CO₂.

For the indicated experiments, cells were exposed to replication stress or treated with specific kinase inhibitors for 22 h prior to the start of the DNA-labeling protocol. Replication stress was induced using 50 µM hydroxyurea (HU, Sigma-Aldrich, H8627) or 0.05 µg/mL aphidicolin (APH, Sigma-Aldrich, A0781). ATM was inhibited using 3 µM KU-600190 (MedChemExpress, HY-1206). ATR inhibition was performed using 5 µM VE-821 (MedChemExpress, HY-14731), 50 nM VE-822(MedChemExpress, HY-13902), or 0.4 µM AZD6738(MedChemExpress, HY-19323). CHK1 was inhibited using 300 nM LY2603618 (MedChemExpress, HY-14720) or 1 µM CCT245737(MedChemExpress, HY-18958), and WEE1 was inhibited using 4 µM MK-1775 (MedChemExpress, HY-10993). CDK2 and CDC7 were inhibited using 8 nM INX-315( MedChemExpress, HY-162001),15 nM PHA-767491(MedChemExpress, HY-13461), respectively.

For experiments involving auxin-inducible degron (AID)-tagged ATR in DT40 cells, indole-3-acetic acid (IAA; I5148 Sigma-Aldrich) was freshly prepared for each experiment as a 0.5 M stock solution in ethanol and added to the culture medium at a final concentration of 0.5 mM for 2 h prior to the start of the DNA-labeling protocol.

All treatments were maintained in the medium throughout the subsequent IdU (Sigma-Aldrich,I7125) and CldU(Sigma-Aldrich,C6891) labeling pulses during DNA combing and was removed during the 1.5-h label-free growth period.

### DNA combing analysis

DNA combing analysis was performed as previously described^48^, with minor modifications. Briefly, unsynchronized cells were pulse-labeled with 25 µM IdU for 30 min, followed by 600 µM CldU for an additional 30 min. The higher concentration of CldU was used to efficiently compete with residual IdU during the second labeling pulse, thereby allowing clear discrimination between the two sequential labeling periods. Cells were then cultured for 1.5 h in label-free medium.

Cells were harvested and embedded in 2% low-melting-point agarose plugs. The plugs were incubated for 48 h at 50°C in digestion buffer containing 1% *N*-lauroylsarcosine, 0.2% sodium deoxycholate, 10 mM Tris-HCl (pH 7.5), 100 mM EDTA, and 1 mg/mL proteinase K. Following digestion, the plugs were incubated overnight in TE ×50 (Tris-EDTA) and stored at 4°C.

Prior to DNA combing, the plugs were washed three times with TE ×1 and once with 50 mM MES (pH 6.0). The plugs were then dissolved in 50 mM MES (pH 6.0) containing 100 mM NaCl. DNA fibers were combed onto Genomic Vision combing slides (CliniSciences, COV-002-RUO) and denatured with 1 M NaOH.

For detection, slides were incubated with the following primary antibodies: rat anti-BrdU (Abcam, ab6326; 1:200), mouse anti-BrdU (BD Biosciences, 347580; 1:20), and anti-DNA antibody (DSHB, AB 10805144; 1:20). The corresponding secondary antibodies were goat anti-mouse Alexa Fluor 647 (Jackson ImmunoResearch, 115-605-166; 1:400), goat anti-rat Alexa Fluor 594 (Thermo Fisher Scientific, A-11007; 1:400), and anti-mouse IgG2a Alexa Fluor 488 (Thermo Fisher Scientific, A-21131; 1:500).

Slides were imaged using a fluorescence microscope, and DNA fibers were manually analyzed using Olympus cellSens software.

In our experiments, cells were sequentially pulse-labeled with IdU and CldU, detected as red and yellow tracts, respectively, while single-stranded DNA (ssDNA) was stained in green.

#### Fork rate measurement

Replication fork rate was determined by measuring IdU (red) tracts that were preceded by a green DNA segment and followed by a CldU (yellow) tract. Only these green–red–yellow patterns were analyzed to ensure that the DNA fiber was intact and that the measured IdU tract represented DNA synthesized throughout the defined IdU labeling period. The IdU tract length was divided by the labeling time to obtain the fork rate in µm/min. The average fork rate calculated for each slide was subsequently used for ROAT calculations

#### Inter-origin distance

IOD was defined as the distance between the centers of two adjacent replication origins. Origins were identified according to their labeling pattern and could be classified into two configurations depending on their activation time relative to IdU addition. Origins activated before the beginning of IdU labeling displayed a yellow–red–green–red–yellow pattern, with the center of the green region defined as the origin center. In contrast, origins activated during the IdU pulse displayed diverging red tracts followed by yellow tracts (yellow–red–yellow), with the center of the continuous red region defined as the origin center. IOD was measured from the center of one origin to the center of the adjacent origin.

#### ROAT and ΔROAT calculation

We further exploited the spatial information obtained by DNA combing to estimate the relative activation timing of individual origins. This measurement was termed **Relative Origin Activation Timing (ROAT)**. For origins activated before IdU addition, the distance spanning the red–green–red region was measured, whereas for origins activated during the IdU pulse, the total red region spanning the two diverging forks was measured. Because DNA synthesis proceeds bidirectionally from each origin, the measured distance represents the combined progression of two replication forks and was therefore divided by two. This value was then converted from distance to time using the average replication fork rate determined for the corresponding slide. ROAT therefore represents the estimated time of activation of an individual origin relative to the start of CldU labeling.

To determine the temporal coordination between two adjacent origins, **ΔROAT** was calculated as the absolute difference between their individual ROAT values:

### ΔROAT = |ROAT₁ − ROAT₂|

Thus, ΔROAT represents the estimated time interval, in minutes, between the activation of two adjacent replication origins. A small ΔROAT indicates closely coordinated origin activation, whereas a larger ΔROAT indicates a greater temporal separation between the activation of the two adjacent origins.

### FACS sorting of early– and late-S-phase cells

For experiments comparing early and late S phase, cells were harvested following the DNA-labelling protocol and were fixed in ethanol in preparation for fluorescence-activated cell sorting (FACS). 600,000 cells from the first third of S phase (Early), or last third of S phase (Late) were collected based on propidium Iodide(PI, Sigma-Aldrich P4170) stains. Following sorting, cells from each fraction were wee mixed with 600,000 unlabeled cells and then embedded in 2% low-melting-point agarose plugs and subsequently processed for DNA combing as described above.

### Flow Cytometry measurements

Cells were fixed in 1% paraformaldehyde in PBS for 20 min at room temperature, washed with PBS and permeabilized in 0.1% saponin for 20 min at 4°C. Flow-cytometric measurements were acquired using a BD LSRFortessaTM Cell **Analyzer** with data being captured and processed using the BD FACSDiva Software. Analysis was conducted using FCS Express 7. Cellular DNA content was measured using DAPI (Sigma, 1 µg/mL in PBS). Cells were filtered through a 35 μm mesh into a new 5 ml polystyrene tube (BD). Cell-cycle gates were manually defined for G1 peaks, and sample’s DAPI signal was normalised by dividing all values by that of the G1 peak.

### Transfection and generation of stable cell lines

HEK293 cells were transfected using TransIT®-LT1 Transfection Reagent(Mirus Bio, MIR 2300), whereas MEFs were transfected by electroporation using the NEPA21 Super Electroporator (program 6). Stable cell lines were generated by selection with G418(Sigma-Aldrich, A1720) at 0.5 mg/mL for HEK293 cells and 0.4 mg/mL for MEFs.

Both cell lines were independently transfected with plasmids encoding N-terminal c-Myc-tagged human *SMC1A*: wild-type *SMC1A* (pCDNA3 5′ cMyc hSMC1 WT; Addgene plasmid #32363), pCDNA3 5′ cMyc hSMC1-S957A/S966A (Addgene plasmid #32372; hereafter referred to as *SMC1A* **-AA**), or two additional constructs generated by GenScript from the WT *SMC1A* plasmid by site-directed mutagenesis, encoding h *SMC1A* –S966E or the double phosphomimetic mutant h *SMC1A* – S957D/S966E (hereafter referred to as *SMC1A* **-DE**). Stable cell lines expressing WT human ATR (pcDNA3-hATR WT; Addgene plasmid #31611), encoding N-terminal FLAG-tagged human ATR, were also generated. The *SMC1A* and ATR expression vectors contain a neomycin-resistance selectable marker, enabling selection with G418. Stable cell lines expressing each construct individually were established and maintained under the corresponding G418 selection conditions.

For experiments requiring simultaneous expression of *SMC1A* –S966E and ATR, the stable *SMC1A* –S966E cell line was additionally transiently transfected with the WT ATR plasmid (pcDNA3-hATR WT; Addgene plasmid #31611).

### Co-immunoprecipitation (Co-IP) and immunoblotting

FLAG-tagged ATR was extracted from a stable HEK293 cell line expressing FLAG-ATR. Cells were washed with PBS and lysed in lysis buffer containing 25 mM HEPES (pH 7.85), 500 mM NaCl, 0.5% NP-40, and 10% glycerol. Prior to the interaction assay, the FLAG-ATR-containing lysate was diluted with NaCl-free lysis buffer of otherwise identical composition to reduce the NaCl concentration to 250 mM.

Myc-tagged wild-type *SMC1A* (*SMC1A* –WT) and the phosphomimetic *SMC1A* – S966E mutant were synthesized in vitro using the TnT® Quick Coupled Transcription/Translation System (Promega, L2081) according to the manufacturer’s instructions. The in vitro-translated Myc-SMC1A-WT or Myc-SMC1A-S966E reaction was combined with the FLAG-ATR-containing cell lysate and incubated for 3 h at 4°C with gentle rotation to allow complex formation.

Following the interaction step, the mixtures were further diluted with NaCl-free lysis buffer to a final NaCl concentration of 150 mM. FLAG-ATR was immunoprecipitated using 5–6 µg of mouse monoclonal anti-FLAG M2 antibody (Sigma-Aldrich, F1804) per reaction. Samples were incubated with the antibody for 3 h at 4°C with gentle rotation, followed by the addition of Protein G beads(Cell Signaling Technology, #70024), and incubation for an additional 1.5 h at 4°C. The beads were subsequently washed with lysis buffer containing 150 mM NaCl.

Immunoprecipitated proteins were eluted by resuspending the beads in 3× sample buffer supplemented with 100 mM DTT (dithiothreitol; Thermo Scientific, R0862), followed by heating at 95°C for 5 min. Immunoprecipitated samples and 5% input samples were resolved by SDS-PAGE using 7.5% polyacrylamide gels. Proteins were transferred to nitrocellulose membranes (Whatman, Cytiva, Cat. No. 10401380) overnight at 30 V using transfer buffer containing 10% methanol and 0.02% SDS.

Membranes were blocked with 5% non-fat dry milk (Sigma-Aldrich, Cat. No. 1153630500) in TBST and subsequently incubated with primary antibodies diluted in 5% non-fat dry milk in TBST. Myc-tagged SMC1A was detected using anti-Myc-tag (9B11) antibody (Cell Signaling Technology, 5698S; 1:750), whereas ATR was detected using anti-ATR antibody (Santa Cruz Biotechnology, sc-515173; 1:500).

### Correlation and Linear Regression Analysis

The relationship between fork rate (FR), Inter Origin Distance (IOD) or Fork symmetry with ΔROAT was assessed using Pearson’s correlation analysis, and the Pearson correlation coefficient (*r*) and corresponding two-sided *p*-value were calculated using the cor.test function in R. Linear regression lines were fitted using ordinary least-squares regression (lm), with ΔROAT as the dependent variable and fork rate as the independent variable. The shaded areas surrounding the fitted regression lines represent 95% confidence intervals, calculated using the default confidence level of the geom_smooth function in the ggplot2 package.

### Super resolution microscopy

Genome replication labeling and visualization: MEF cells were seeded on high-precision #1.5H glass coverslips (170 ± 5 µm thickness) and allowed to attach onto glass before treatment. Cells were pulse-labelled with 10 µM of EdU (Cat.No.: 7845.1, Click-IT-EdU cell proliferation assay, Carl Roth, Karlsruhe, Germany), for 10 min before washing with PBS 1× and fixing with 3.7% formaldehyde in PBS 1× for 10 min. The cells were washed thrice for 5 min each with PBS 1×. All the immunostaining was performed inside a dark, humidified chamber at room temperature unless otherwise mentioned. After washing, the cells were permeabilized with 0.5% Triton X-100 (Carl Roth, Karlsruhe, Germany) in PBS 1× for 10 min. The cells were washed thrice for 5 min each with a washing buffer (0.05% Tween in PBS 1×) and blocked with 4% BSA in PBS 1× for 30 min. For the detection of EdU, cells were incubated in Click-IT cocktail mix 100 mM Tris-HCl, pH 8.5, 10 mM CuSO4, 1 µM 5-TAMRA-Azide (Cat.No.: CLK-FA008-1, Jena Bioscience, Jena, Germany), and 100 mM ascorbic acid diluted in water for 30 min (Salic and Mitchison 2008). All the cells were stained with 10 mg/mL DAPI (4′,6-diamidino-2-phenylindole, Cat.No.: D9542, Sigma-Aldrich Chemie GmbH, Steinheim, Germany) for 10 min and mounted on Vectashield (Cat.No.: VEC-H-1000, Vector Laboratories Inc., Burlingame, CA, USA). All the coverslips were sealed with transparent nail polish and air-dried in the dark.

Super-resolution imaging was performed on a Zeiss LSM 900 laser-scanning confocal microscope equipped with an Airyscan 2 detector (32-element GaAsP-PMT array), mounted on an Axio Observer 7 inverted stand and operated with ZEN blue 3.9 (Carl Zeiss Microscopy, Jena, Germany). Images were acquired with a C Plan-Apochromat 63×/1.40 Oil DIC M27 objective using immersion oil of refractive index 1.518. Excitation was provided by 405 nm (DAPI) and 561 nm (EdU, TMR) diode lasers at 11.4 % and 6.0 % relative transmission, respectively, and directed to the sample through an MBS 405/561/640 main beam splitter. Emission was collected in the ranges 400–605 nm (DAPI) and 540–700 nm (TMR), with the two channels recorded in frame-wise sequential mode to avoid bleed-through. Acquisition was performed in Airyscan super-resolution (SR) mode with the pinhole set to 5.0 AU (TMR and Alexa Fluor 488) and 5.8 AU (DAPI). Images of 958 × 958 pixels were acquired at 3× optical zoom, giving a lateral pixel size of 35.3 nm and a field of view of 33.8 × 33.8 µm, which satisfies Nyquist sampling for this mode. Z-stacks of 43 optical sections were recorded at a step size of 140 nm, covering approximately 6 µm and spanning the full nuclear volume. Raw Airyscan datasets were processed in ZEN blue 3.9 using Airyscan joint deconvolution (jDCV) with 20 iterations, a quality threshold of 0.005 and the “Dense” sample-structure setting, applied identically to all channels and all experimental conditions. Both airyscan and airyscan with joint deconvolution images were exported for further analysis.

### Image analysis

All image analysis was performed in Fiji/ImageJ using the 3D ImageJ Suite as earlier ^49^ with minor modifications. One nucleus was quantified per image, and nuclei were classified as early-or late-S-phase based on the spatial replication labelling pattern. Custom ImageJ macros were used to batch-process the datasets, and identical parameters were applied to every image across all conditions. Airyscan– and airyscan with joint deconvolution-processed. CZI files were exported as TIFF hyperstacks, each channel was independently scaled to its own stack minimum and maximum and converted to 8-bit, and the stacks were then split into single-channel files (C1, EdU-TMR; C2, DAPI). Nuclear masks were generated from the DAPI channel. The EdU channel was restricted to the nuclear volume by a voxel-wise minimum operation with this mask

Replication (nano)foci were segmented from the nucleus-masked EdU-TMR channels (**Figure S4**). Stacks were smoothed with a 3D mean filter (radius 1 px) and intensity-normalized across the stack. Local maxima were detected plane-by-plane (Find Maxima, noise tolerance 1, output “Single Points”), and an intensity-weighted seed image was generated by a voxel-wise minimum between the normalized stack and the maxima map (https://imagej.net/ij/macros/FindStackMaxima.txt). Seeded 3D segmentation was then performed with 3D Spot Segmentation (seed threshold 50; local threshold estimated by Gaussian fit; radii 2, 4 and 6 px with a weight of 0.50; maximum radius 5 px; SD value 1.40; classical spot segmentation with watershed), restricting accepted objects to a volume of 50–300 voxels (≈ 0.009–0.052 µm³). Foci remaining in contact after segmentation were separated by 3D Watershed Split (radius 2 px) using the seed image as marker. Segmented (nano)RFI were assigned to their parent nucleus with 3D Numbering, using the DAPI-derived mask as the region of interest and the split label image as the counted objects, returning for each nucleus the number of replication foci, their cumulative volume and the fraction of the nuclear volume they occupy. Furthermore, the number of nano RFi within individual RFi was quantified using 3D numbering with segmented nano RFi as objects within individual RFi as the volume of interest.

## Acknowledgments

The authors thank the Simon lab members for fruitful discussions. We are thanking Prof. Michael B Kastan for sending us the *Smc1a* – S957A, S966A MEF cells, Prof. Noel Francis Lowndes for sending the AID-ATR DT40 cells and Prof. Sara Buonomo for the HEK293 Rif1 KO cells.

## Data availability

The data that support the findings of this study are available from the corresponding author upon request.

## Author contributions

I.S. conceived the study; J.K., I.S. and S.E. designed the research. J.K. performed most of the experiments in Figures 1-3. S.K.P., M.K.P., and M.A. performed super-resolution imaging and analysis; A.G. and L.D. performed cell-cycle analysis and T-cell experiments. I.S. and M.C.C. supervised the research and acquired funding; I.S. wrote the manuscript with input from all authors.

