## supplementary figures 1,2,3,4 for "Constitutive ATR Signaling Enforces Domain-Wide Origin Synchrony"

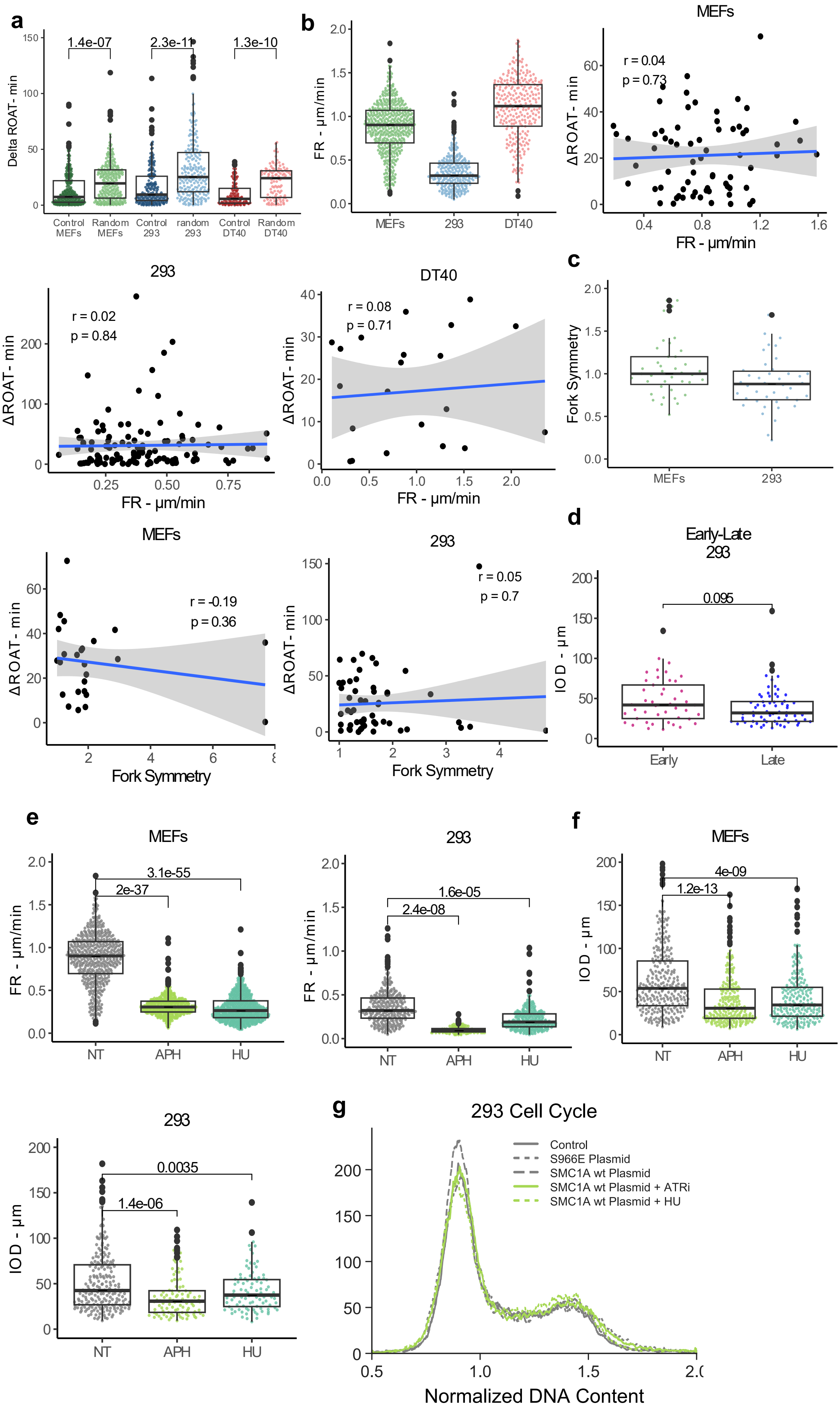
a
Delta ROAT- min
1.4e-07
2.3e-11
1.3e-10
Control MEFs
Random MEFs
Control 293
random 293
Control DT40
Random DT40
b
FR - µm/min
MEFs
293
DT40
MEFs
r = 0.04
p = 0.73
ΔROAT - min
FR - µm/min
293
r = 0.02
p = 0.84
ΔROAT - min
FR - µm/min
DT40
r = 0.08
p = 0.71
ΔROAT - min
FR - µm/min
c
Fork Symmetry
MEFs
293
MEFs
r = -0.19
p = 0.36
ΔROAT - min
Fork Symmetry
293
r = 0.05
p = 0.7
ΔROAT - min
Fork Symmetry
d
Early-Late
293
0.095
IOD - µm
Early
Late
e
MEFs
3.1e-55
2e-37
FR - µm/min
NT
APH
HU
293
1.6e-05
2.4e-08
FR - µm/min
NT
APH
HU
f
MEFs
4e-09
1.2e-13
IOD - µm
NT
APH
HU
293
0.0035
1.4e-06
IOD - µm
NT
APH
HU
g
293 Cell Cycle
Control
S966E Plasmid
SMC1A wt Plasmid
SMC1A wt Plasmid + ATRi
SMC1A wt Plasmid + HU
Normalized DNA Content

**Supplementary Figure 1. FR and IOD and their association to ΔROAT.**

(a)ΔROAT between adjacent replication origins in MEFs, HEK293, and DT40 cells compared with ΔROAT values obtained from randomly paired origins (Random). **(b)** Fork rate (FR) in MEFs, HEK293, and DT40 cells (left), and correlation between FR and ΔROAT in each cell line (right). Each point represents an individual measurement. For the correlation analyses, regression lines and corresponding confidence intervals are shown. Pearson correlation coefficients (r) and corresponding p values are indicated in each graph. **(c)** Fork symmetry in MEFs and HEK293 cells (upper, left), and correlation between fork symmetry and ΔROAT in each cell line (lower, right). Fork symmetry represents the ratio between the lengths of the two replication forks extending bidirectionally from the same origin. Each point represents an individual measurement. The Correlation analyses were performed as in **a**. **(d)** Inter-origin distance (IOD) in early- and late-S-phase HEK293 cells. Fork rate **(e)** and IOD **(f)** in untreated (NT) or treated with aphidicolin (APH) or hydroxyurea (HU) in MEFs(left) and 293 cells(right). Each point represents an individual measured replication fork. P values for the indicated comparisons (two-tailed t-test) are shown above the plots. Details about the inhibitors are in the Methods section **(g)** Cell-cycle distribution of HEK293 under control conditions, following expression of WT or S966E SMC1A, and following ATR inhibition or HU treatment in WT SMC1A - expressing cells. DNA content was assessed by DAPI staining, and the distributions are shown as normalized DAPI intensity versus cell count.

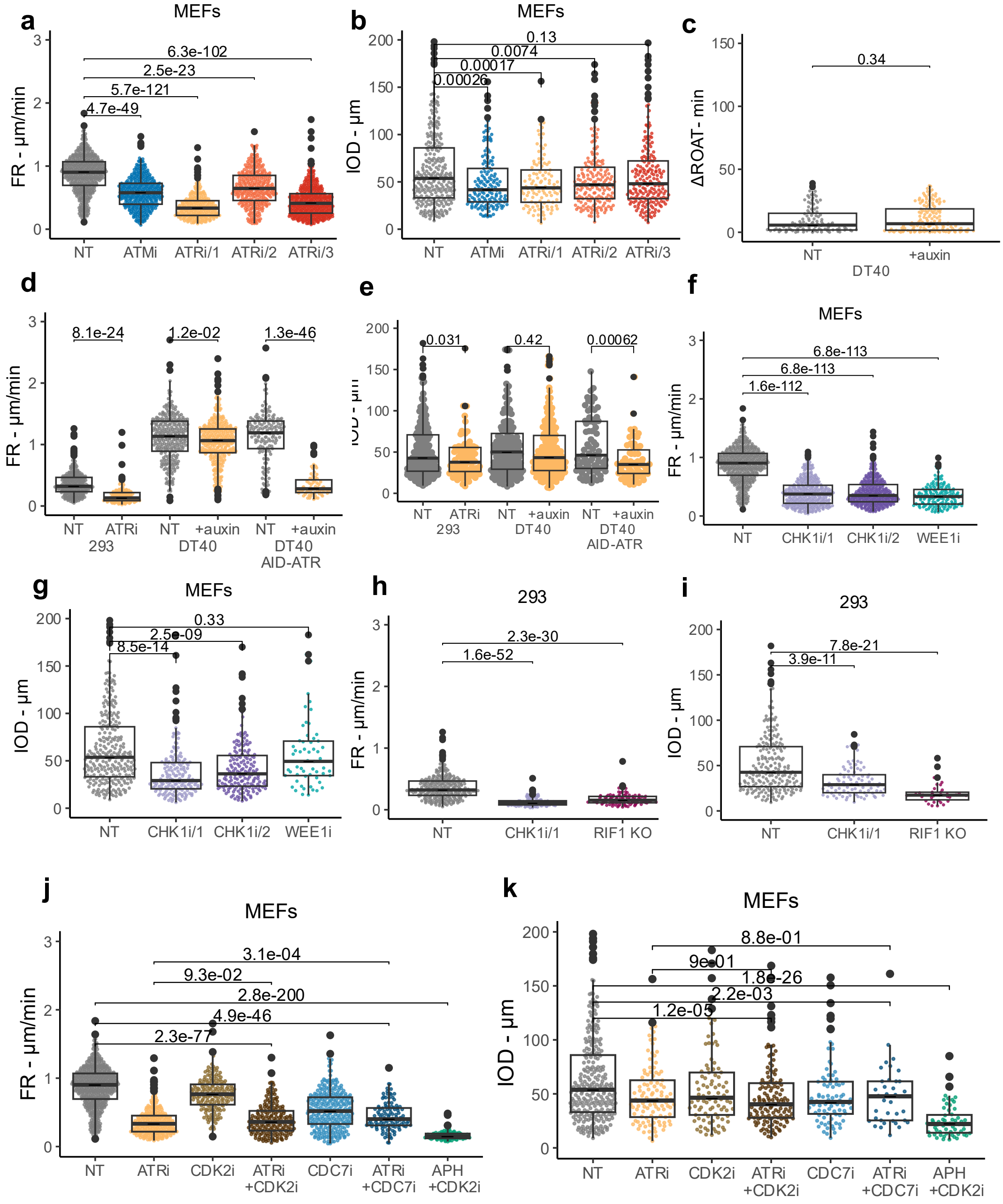


**Supplementary Figure 2. Effects of ATR–CHK1 pathway inhibition on replication fork rate and inter-origin distance.**

**(a,b)** Replication fork rate (FR) **(a)** and inter-origin distance (IOD) **(b)** in untreated (NT) MEFs and following inhibition of ATM (ATMi) or ATR using three ATR inhibitor conditions. **(c)** ΔROAT between adjacent replication origins in DT40 WT cells in the absence (NT) or presence of auxin-induced ATR depletion. **(d, e)** FR **(d)** and IOD **(e)** in HEK293 cells following ATR inhibition (VE-821), and in DT40 WT and AID-ATR cells in the absence or presence of auxin. **(f, g)** FR **(f)** and IOD **(g)** in untreated (NT) MEFs, following treatment with two CHK1 inhibitors, or following treatment with one WEE1 inhibitor. **(h, i)** FR **(h)** and IOD **(i)** in untreated (NT) HEK293 cells, following CHK1 inhibition or in RIF1 KO cells. **(j, k)** FR **(j)** and IOD **(k)** in MEFs following ATR inhibition alone or in combination with CDK2 or CDC7 inhibition, plus MEFs treated with APH combined with CDK2i. *P* values for the indicated comparisons (two-tailed t-test) are shown above the plots. Details about the inhibitors are in the Methods section.

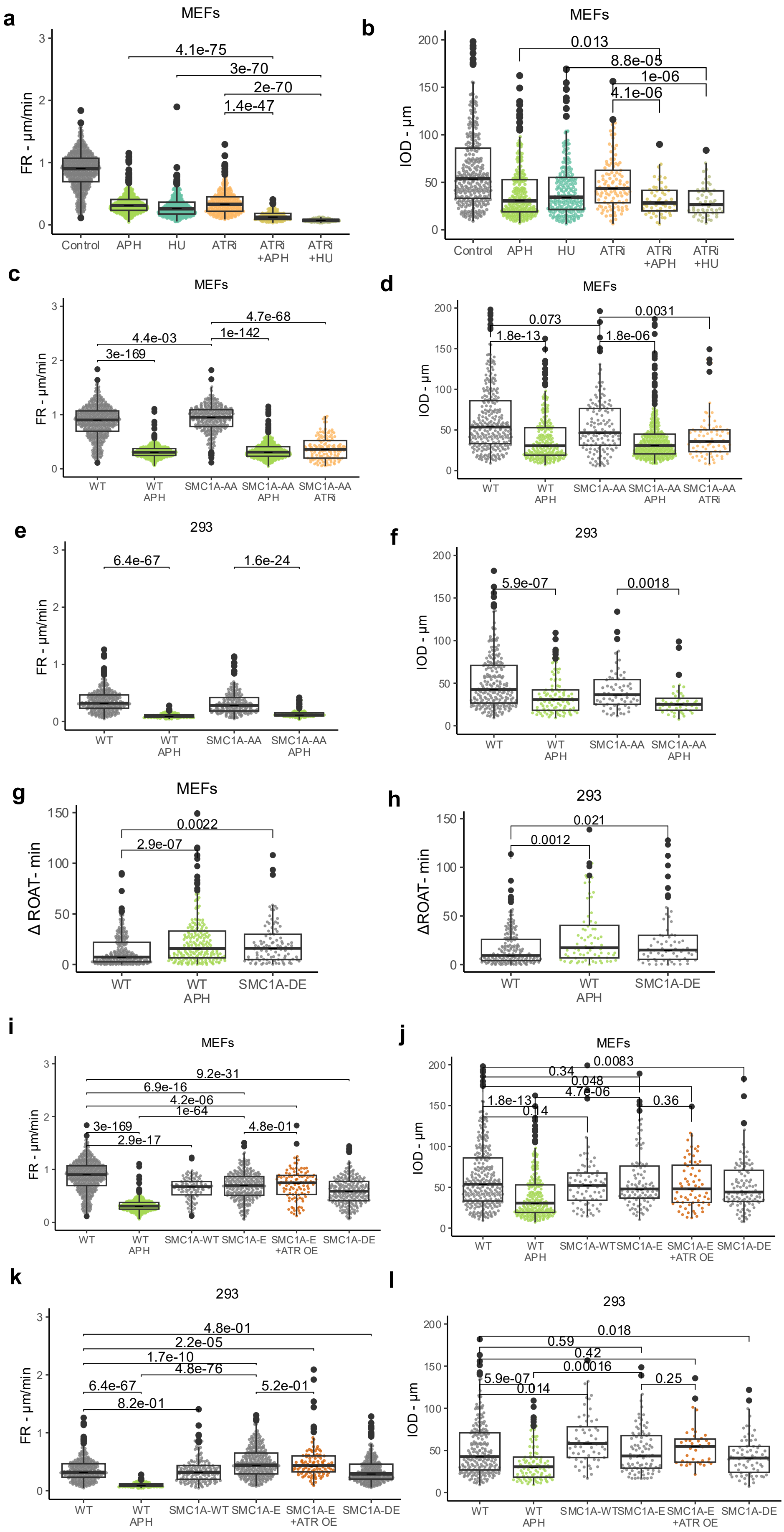

a
MEFs
FR - µm/min
4.1e-75
3e-70
2e-70
1.4e-47
Control
APH
HU
ATRi
ATRi +APH
ATRi +HU
b
MEFs
IOD - µm
0.013
8.8e-05
1e-06
4.1e-06
Control
APH
HU
ATRi
ATRi +APH
ATRi +HU
c
MEFs
FR - µm/min
4.7e-68
4.4e-03
1e-142
3e-169
WT
WT APH
SMC1A-AA
SMC1A-AA APH
SMC1A-AA ATRi
d
MEFs
IOD - µm
0.073
0.0031
1.8e-13
1.8e-06
WT
WT APH
SMC1A-AA
SMC1A-AA APH
SMC1A-AA ATRi
e
293
FR - µm/min
6.4e-67
1.6e-24
WT
WT APH
SMC1A-AA
SMC1A-AA APH
f
293
IOD - µm
5.9e-07
0.0018
WT
WT APH
SMC1A-AA
SMC1A-AA APH
g
MEFs
Δ ROAT- min
0.0022
2.9e-07
WT
WT APH
SMC1A-DE
h
293
ΔROAT- min
0.021
0.0012
WT
WT APH
SMC1A-DE
i
MEFs
FR - µm/min
9.2e-31
6.9e-16
4.2e-06
1e-64
3e-169
4.8e-01
2.9e-17
WT
WT APH
SMC1A-WT
SMC1A-E
SMC1A-E +ATR OE
SMC1A-DE
j
MEFs
IOD - µm
0.0083
0.34
0.048
4.7e-06
1.8e-13
0.36
0.14
WT
WT APH
SMC1A-WT
SMC1A-E
SMC1A-E +ATR OE
SMC1A-DE
k
293
FR - µm/min
4.8e-01
2.2e-05
1.7e-10
4.8e-76
6.4e-67
5.2e-01
8.2e-01
WT
WT APH
SMC1A-WT
SMC1A-E
SMC1A-E +ATR OE
SMC1A-DE
l
293
IOD - µm
0.018
0.59
0.42
0.00016
5.9e-07
0.25
0.014
WT
WT APH
SMC1A-WT
SMC1A-E
SMC1A-E +ATR OE
SMC1A-DE

**Supplementary Figure 3. SMC1A phosphorylation contributes to replication-stress-induced changes in replication dynamics and origin coordination.**

**(a,b)** FR **(a)** and IOD **(b)** in WT MEFs under control conditions, following replication stress induced by aphidicolin (APH) or hydroxyurea (HU), and following ATR inhibition (ATRi) alone or in combination with APH or HU. **(c–f)** Fork rate (**c,e**) and IOD (**d,f**) in WT and SMC1A-AA MEFs (c-d) or HEK293 cells (e-f) under untreated conditions or following APH treatment or ATRi(VE-821) in MEFs. **(g–h)** ΔROAT in WT cells, APH-treated WT cells, and cells expressing the phosphomimetic SMC1A -DE mutant in MEFs (**g**) and HEK293 cells (**h**). **(i–l)** FR **(i,k)** and IOD **(j,l)** in WT, APH-treated and cells expressing WT SMC1A, the phosphomimetic SMC1A-DE and SMC1A-E mutants, with or without ATR overexpression (ATR OE) in MEFs **(i-j)** and HEK293 cells **(k-l).** *P* values for the indicated comparisons (two-tailed t-test) are shown above the plots. Details about the inhibitors are in the Methods section.

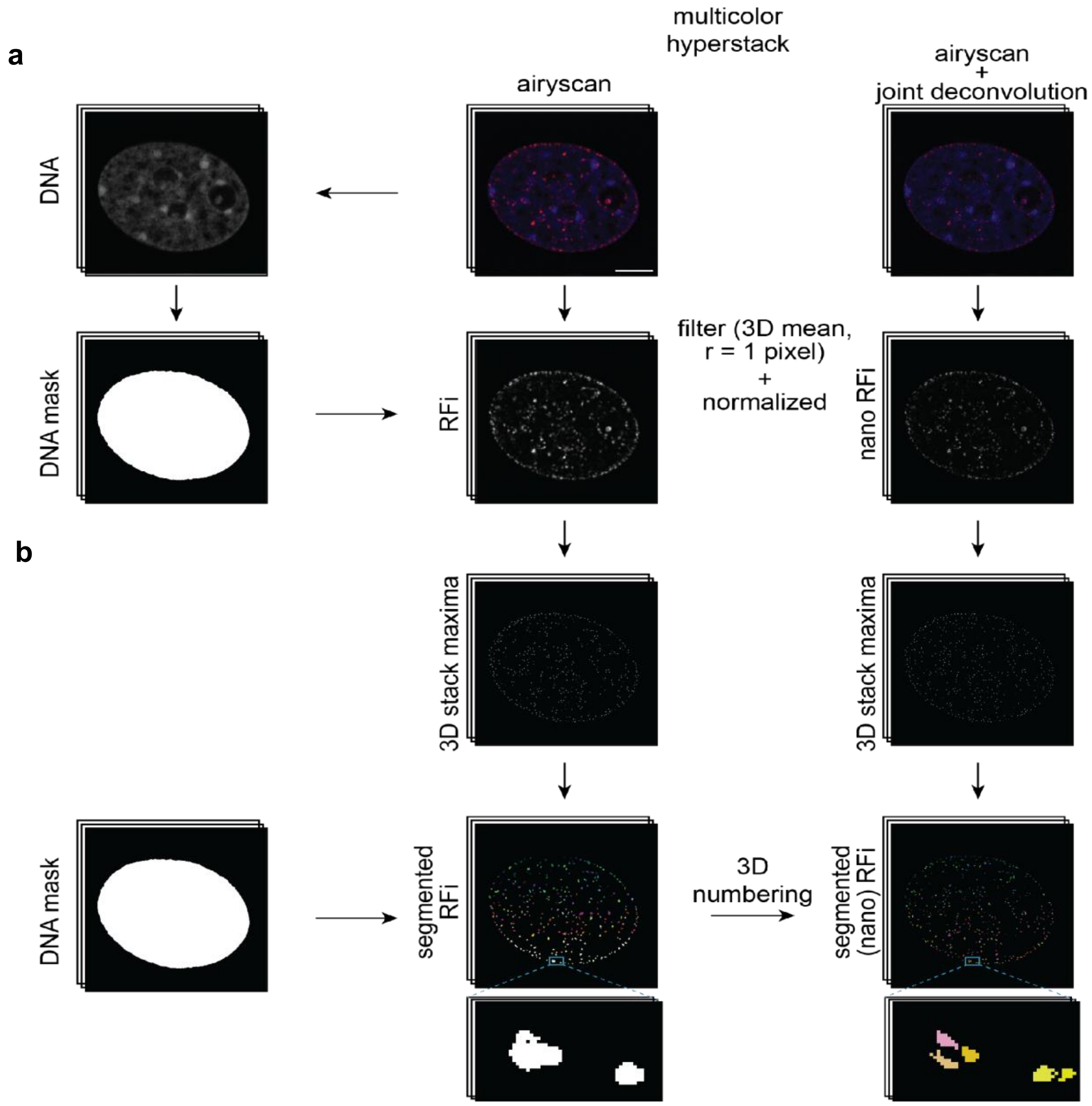


**Supplementary Figure 4: Image analysis pipeline for correlative microscopy.**

**(a)** Processed 8-bit multicolor airyscan & airyscan+joint deconvolution images of the same cells were processed in parallel. DNA (DAPI) channel was used to segment nuclei and define the region of interest. A minimum operation between the DNA mask and (nano) RFi channels was performed. **(b)** Filtered and normalized images were used to find 3D stack maxima, followed by a voxel-wise minimum between the normalized stack and the maxima to get the intensity-weighted seed. Seeded 3D segmentation was then performed with 3D Spot Segmentation, followed by a watershed operation. The zoomed images below show the segmented (nano) RFi at both resolutions.
